# A single-cell and multi-platform spatial atlas of the human pancreas resolves a transformation-associated epithelial axis and a recurrent boundary-organized tumour microenvironment

**DOI:** 10.64898/2026.07.31.742073

**Authors:** Kaili Liu, Hua Zhong, Rahhul S. Elangovan, Yuanhong Sun, Lin Wang, Abigae P. Williams, Trisha I. Valerio, Coline Furrer, Jacob P. Adams, Ashley R. Hoover, Wei R. Chen

## Abstract

Most pancreatic ductal adenocarcinoma (PDAC) single-cell and spatial studies analyze one cohort or platform, obscuring recurrent biology. We assembled a human pancreas single-cell and single-nucleus reference of 1,186,130 cells from 19 studies and interpreted 176 Visium sections comprising 458,877 spots across non-diseased pancreas, chronic pancreatitis, PanIN, IPMN, primary PDAC and metastasis. Marker-supported labels were used after two RNA-based copy-number callers failed known-diploid controls. BANKSY domains, two reference-mapping methods and sample-level analyses resolved a cross-sectional epithelial axis extending from acinar-rich to malignant tissue. Five trajectory algorithms recovered similar ordering on a shared embedding; their consensus was interpreted as transformation-associated, not temporal or clonal. Malignant regions were globally segregated from fibroblast and myeloid compartments. Signed-distance analysis refined this pattern into a malignant core, a CAF/myeloid surround beginning at the tumour boundary and a more distal lymphoid compartment. Candidate extracellular-matrix communication, led by COLLAGEN, LAMININ and FN1, concentrated at the interface.

Changes were reproduced in six patient-matched Normal-tumour pairs using exact patient-level tests. Visium HD resolved the same organization at single-cell resolution and showed that 8-um bins distorted immune-adjacency estimates. Xenium also revealed recurrent neighbourhoods but sample-specific stromal boundaries. We provide a confound-aware framework for identifying recurrent epithelial and microenvironmental organization in PDAC.

## Introduction

Pancreatic ductal adenocarcinoma remains one of the most lethal common malignancies because it is usually detected late, disseminates early and rarely shows a durable response to systemic treatment^1,2^. Histopathological models place pancreatic intraepithelial neoplasia (PanIN) and intraductal papillary mucinous neoplasm (IPMN) before invasive cancer, while genomic studies describe punctuated and branched acquisition of canonical driver alterations^3–5^. Bulk molecular classifications further separate tumour-intrinsic and stromal phenotypes^6–9^. These frameworks are indispensable, but they do not show how malignant, stromal and immune populations are arranged in tissue.

Single-cell studies have resolved malignant epithelial states, fibroblast diversity, precursor heterogeneity and tumour-associated immune programs^10–15^. Later atlases added microenvironment-dependent plasticity, tumour-infiltrating lymphocyte trajectories, metastatic ecosystems and treatment-associated states^16–20^. Integrated and spatially resolved studies have broadened the common cell-state vocabulary^21–23^. Spatial transcriptomics then supplied the coordinate system needed to examine intact PDAC tissue^24–28^.

The central difficulty is separating recurrent disease biology from cohort and platform structure. Public studies differ in tissue preservation, single-cell versus single-nucleus profiling, 3-prime versus 5-prime chemistry, whole-transcriptome versus targeted assays, and definitions of normal pancreas. Healthy donor pancreas is usually acinar-rich, whereas tumour-adjacent tissue can contain fibrosis, inflammation and field effects. In a pooled dataset, publication, preservation and disease can therefore become nearly collinear. The same concern applies to stromal interpretation. PDAC fibroblasts occupy inflammatory, myofibroblastic, antigen-presenting and LRRC15-positive states, and their position relative to malignant glands is part of their biology^14,29–31^. Fibroblasts and their CXCL12 axis can also restrict lymphocyte access^32^, while the broader PDAC stroma has both tumour-promoting and tumour-restraining features^33,34^.

We therefore integrated the data around explicit evidence levels. A 19-study single-cell and single-nucleus atlas supplied the reference vocabulary. Standard Visium provided the discovery cohort. A preservation-restricted analysis reduced one major technical confound, and six patient-matched Normal-tumour pairs supplied the principal within-patient comparison. Visium HD tested whether spot-level geometry remained visible after cell segmentation. Xenium provided a targeted in-situ test whose panel omissions and annotation failures were retained rather than hidden. Samples or patients, rather than cells or spots, were used as biological replicates for group inference.

Two questions guided the analysis. First, could cross-sectional epithelial states be organized along a reproducible molecular coordinate without assuming that RNA-inferred copy number correctly identified malignancy? Second, did malignant, stromal and immune compartments form a recurrent spatial organization across cohorts, matched patients and measurement resolutions? The resulting atlas supports a transformation-associated epithelial axis and a malignant-core, stromal-surround and lymphoid-exterior architecture, while identifying where those conclusions remain sample- or panel-limited.

## Results

### A 19-study single-cell reference separates recurrent pancreatic cell states

The integrated reference contained 1,186,130 cells and nuclei and resolved 17 broad cell types (Fig. 1a-c). T cells, malignant epithelium, CAFs, myeloid cells, acinar cells and ductal cells were the largest compartments; rare Schwann, gastric-type and tuft populations were retained so that their spatial signal was not forced into broad residual labels. Canonical marker blocks, rather than embedding position alone, supported the final annotation. A 28-state vocabulary was kept for lineage-specific analyses, whereas the 17-type vocabulary was used for spatial mapping.

**Fig. 1.**
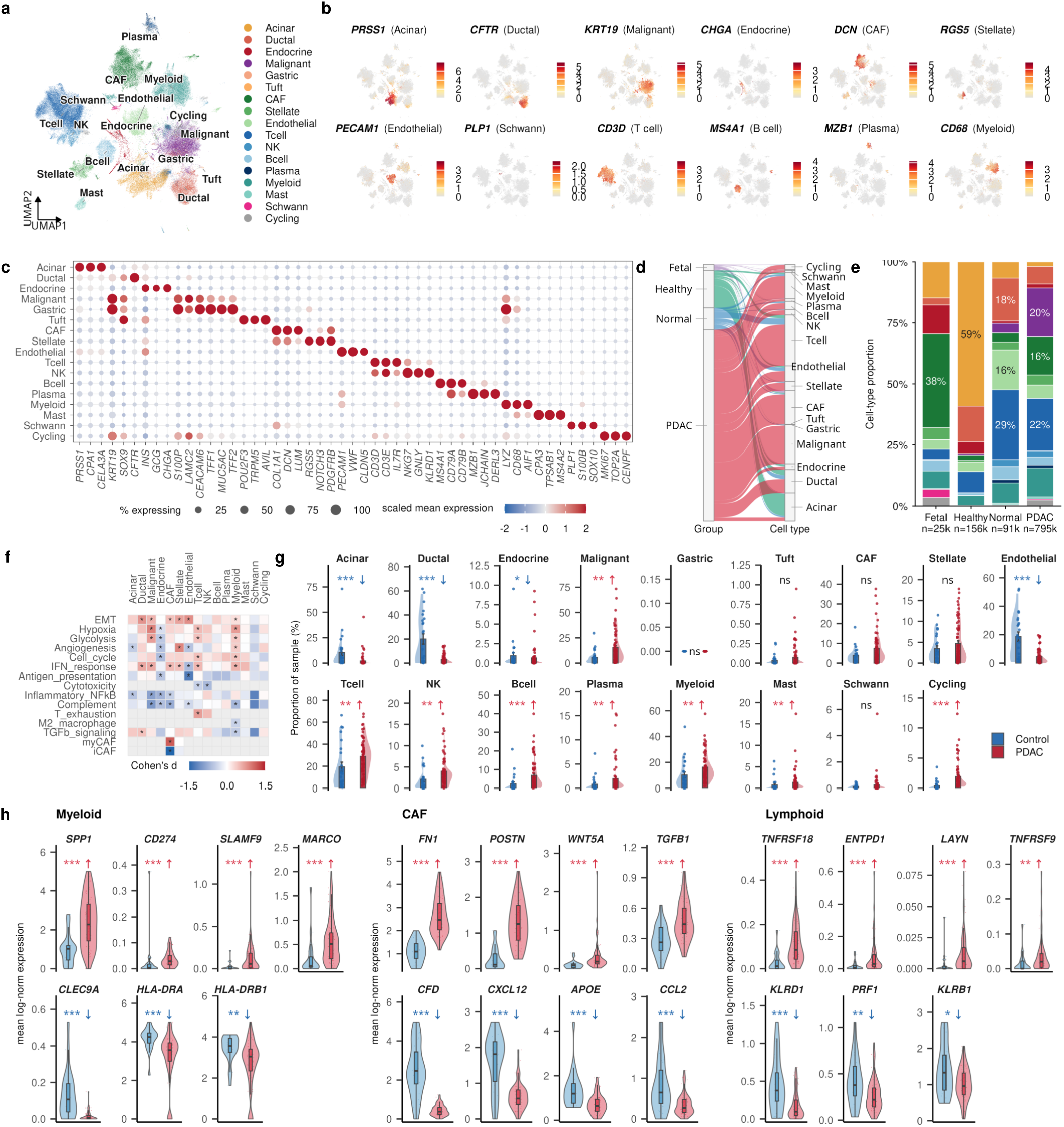
A 19-study single-cell and single-nucleus reference of the human pancreas. **a**, Harmony UMAP of 1,186,130 cells and nuclei colored by 17 broad cell types. **b**, Representative lineage markers on the reference embedding. **c**, Dot plot of 51 canonical markers; dot area denotes percentage expressing and color denotes per-gene z-scored mean expression. **d**, Source group to final cell-type alluvial map. **e**, Cell-type composition of fetal, healthy donor, tumour-adjacent Normal and PDAC groups. **f**, Cell-type-resolved PDAC-Control signature effects from sample-level scores. **g**, Cell-type percentage per sample in PDAC (n=65) and Control (n=24) within five studies containing both arms; two-sided Wilcoxon tests were corrected across 17 types. CAF q=0.057 is not significant. **h**, Sample-level suppressive and pro-immune marker expression in myeloid, CAF and lymphoid compartments using the same study-restricted design. Displayed genes were selected after testing and their P values are descriptive.

RNA-inferred copy number did not provide a reliable malignant-cell anchor. CopyKAT called 99.6% of normal gastric-type epithelial cells and 52% of fetal cells aneuploid, and inferCNV did not separate the relevant malignant and non-malignant epithelial groups above chance. Malignant identity was therefore assigned from epithelial lineage, tumour-associated markers and loss of normal exocrine programs. The copy-number outputs were retained as a negative-control audit rather than used to train or validate downstream labels.

Source composition differed markedly (Fig. 1d,e). Healthy donor pancreas was 59.1% acinar, whereas tumour-adjacent Normal tissue contained more T, ductal and endothelial cells and only 6.7% acinar cells. PDAC contained malignant epithelium, CAF, myeloid and T-cell compartments. These differences supported keeping healthy donor and tumour-adjacent Normal labels separate in descriptive analyses.

The primary PDAC-Control test was restricted to five studies containing both arms after removal of sorted or compositionally non-representative samples. This yielded 65 PDAC and 24 Control samples. Acinar, ductal, endocrine and endothelial fractions decreased, whereas malignant, myeloid, T, NK, B, plasma and cycling-cell fractions increased (Fig. 1f,g). For example, median acinar abundance fell from 4.79% to 0.13% (q=1.37x10^-4), malignant abundance rose from 3.35% to 9.87% (q=0.00194), and myeloid abundance rose from 6.17% to 13.68% (q=0.00449). CAF abundance increased from 3.05% to 4.52% but did not pass correction (q=0.057).

Expression changes indicated functional remodeling rather than immune abundance alone (Fig. 1h). PDAC myeloid cells showed more *SPP1*, *CD274*, *SLAMF9* and *MARCO* and less antigen-presentation signal; CAFs gained *FN1*, *POSTN*, *WNT5A* and *TGFB1*; lymphoid cells gained regulatory or exhaustion-associated genes and lost cytotoxic-associated genes. These markers were selected after testing in the same samples, so their P values are descriptive. cell2location and RCTD agreed most strongly for abundant, spatially coherent compartments, providing a cross-algorithm bridge to spatial data without constituting independent annotation validation.

### Spatial domains define a cross-sectional pancreatic disease axis

The standard-Visium cohort contained 176 sections and 458,877 tissue spots. BANKSY resolved 14 recurrent domains, including acinar, endocrine-islet, malignant mucinous/intestinal, malignant basal/squamous, CAF-desmoplastic, myeloid-TAM, immune-lymphoid/TLS and Schwann- neural territories (Fig. 2a,b). cell2location and RCTD supplied a complementary cell-type view, and their assignments were most concordant in the large acinar, CAF and malignant compartments (Fig. 2c,d). Representative maps showed how the same domain vocabulary was deployed in Normal, chronic pancreatitis, PanIN, primary PDAC and metastasis (Fig. 2e).

**Fig. 2.**
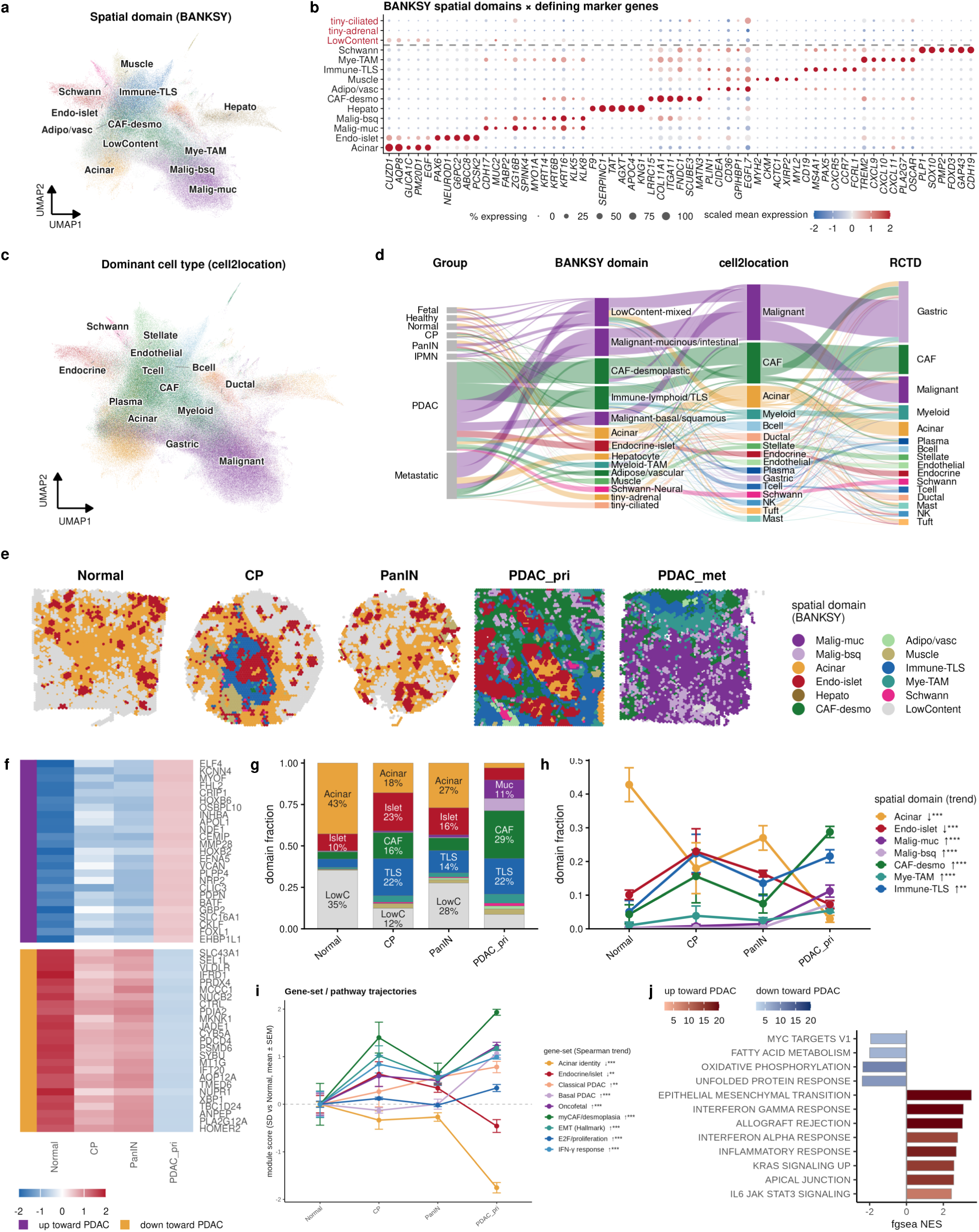
Spatial domains and molecular programs across the pancreatic disease spectrum. The full cohort contains 176 sections and 458,877 spots. **a**, Integrated spatial UMAP colored by 14 BANKSY domains. **b**, Endogenous spatial-marker dot plot. **c**, Dominant cell2location type. **d**, Group, BANKSY domain, dominant cell2location type and RCTD alluvial map. **e**, Representative domain maps from Normal, chronic pancreatitis, PanIN, primary PDAC and metastasis. **f**, Sample-pseudobulk expression of the 50 genes most strongly associated with the ordered FFPE axis. **g**, Mean domain composition. **h**, Per-sample domain abundance and sample-level Spearman tests. **i**, Molecular-program scores relative to the Normal mean. **j**, Hallmark enrichment of the stage-associated gene ranking. Panels **f-j** use 75 FFPE samples; chronic pancreatitis has n=3 and is interpreted cautiously.

Disease state and preservation were partially confounded, so the quantitative stage analysis was restricted to 75 FFPE Normal, chronic-pancreatitis, PanIN and primary-PDAC samples. Sample- pseudobulk genes formed a coherent Normal-to-PDAC expression gradient (Fig. 2f). Acinar- domain abundance fell from 42.8% in Normal to 3.0% in PDAC, whereas CAF-desmoplastic tissue rose from 4.3% to 28.7% and immune-lymphoid/TLS tissue from 5.0% to 21.6% (Fig. 2g). Sample-level trends were strongest for loss of acinar tissue (rho=-0.77, FDR=1.06x10^-14) and gain of malignant basal/squamous (rho=0.69), CAF-desmoplastic (rho=0.64), malignant mucinous/intestinal (rho=0.57) and myeloid-TAM domains (rho=0.48; Fig. 2h).

Molecular programs changed in parallel (Fig. 2i). PDAC gained myCAF/desmoplasia, oncofetal, EMT, basal-PDAC, interferon-gamma and classical-PDAC scores while losing acinar and islet identity. Chronic pancreatitis already showed strong desmoplasia and EMT, indicating that these programs are not cancer specific. The chronic-pancreatitis group contained only three samples and was not used to infer a finely resolved lesion sequence. Hallmark analysis identified EMT as the strongest positive pathway (NES=3.53), followed by interferon and inflammatory programs; secretory and metabolic programs associated with differentiated exocrine tissue declined (Fig. 2j). These are cross-sectional sample-level associations, not longitudinal transitions.

### A transformation-associated epithelial axis is robust to trajectory algorithm

We selected 83,384 epithelial-enriched FFPE spots from 84 samples and applied diffusion pseudotime, Slingshot, VIA, Palantir and Monocle3 to the same Harmony representation. All methods were rooted in acinar epithelium. Palantir ranked highest among individual tools, but the rank-mean consensus was most stable across pathological-group recovery, cross-tool agreement, spatial autocorrelation, malignant-fraction association, endpoint composition and bootstrap stability. Because all methods shared one embedding, root and upstream labels, their agreement measures algorithmic robustness rather than independent lineage evidence.

The consensus formed a continuous UMAP gradient (Fig. 3a), increased across malignant and desmoplastic domains (Fig. 3b), and correlated with deconvolved malignant fraction (spot-level rho=0.72; Fig. 3c). Tissue maps showed broad low scores in non-diseased pancreas, focal higher-score regions in chronic pancreatitis and PanIN, and extensive high-score territories in primary and metastatic disease (Fig. 3d). The malignant-fraction comparison is not independent because both quantities reuse the reference, whereas the spatial smoothness check addresses a different failure mode.

**Fig. 3.**
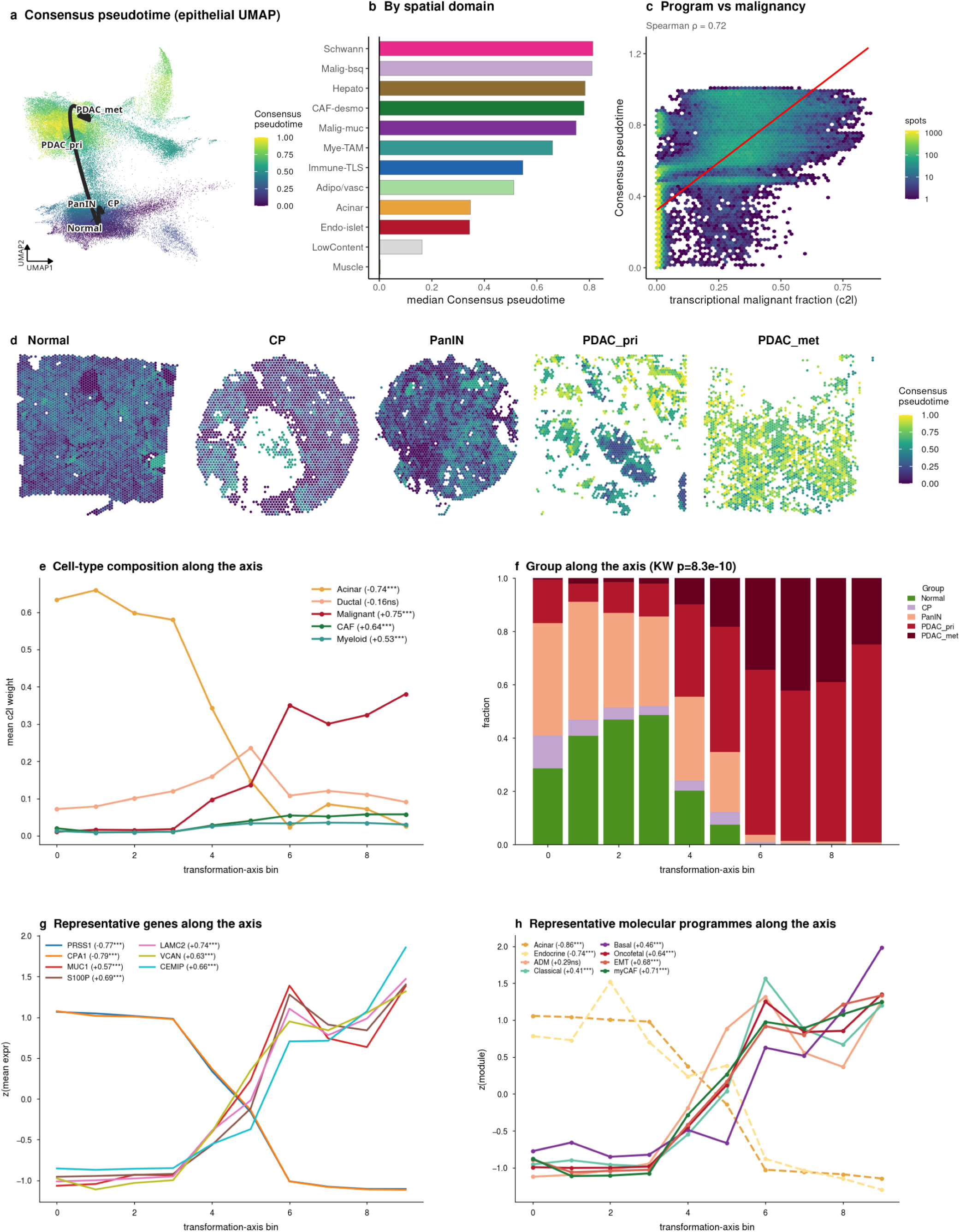
A cross-sectional transformation-associated epithelial axis. The analysis contains 83,384 epithelial-enriched FFPE spots from 84 samples. **a**, Rank-mean consensus of five trajectory methods on a common Harmony embedding rooted in acinar epithelium. **b**, Median consensus by BANKSY domain. **c**, Consensus versus cell2location malignant fraction (spot-level rho=0.72; descriptive because both reuse the reference). **d**, Representative tissue maps. **e**, Cell-type composition along the axis with sample-level correlations. **f**, Per-sample score by pathological group (Kruskal-Wallis P=8.31x10^-10). **g**, Representative endogenous genes. **h**, Molecular-program associations; ADM/metaplasia does not pass correction (q=0.052). The consensus is an ordering of cross-sectional similarity, not a temporal or clonal trajectory.

Cellular composition changed continuously along the axis (Fig. 3e). Sample-level acinar abundance declined (rho=-0.74), while malignant (rho=0.75), CAF (rho=0.64) and myeloid abundance (rho=0.53) increased. Ductal and endocrine abundance were not significant. Per-sample scores differed across pathological groups (Kruskal-Wallis P=8.31x10^-10; Fig. 3f). Acinar genes *PRSS1* and *CPA1* declined, whereas *MUC1*, *S100P*, *LAMC2*, *VCAN* and *CEMIP* increased (Fig. 3g). Acinar and islet programs decreased and myCAF/desmoplasia, EMT, oncofetal, classical-PDAC and basal-PDAC programs increased (Fig. 3h); ADM/metaplasia did not pass correction (q=0.052).

Leave-one-study-out and unintegrated analyses preserved the broad ordering. We therefore call the score a transformation-associated epithelial axis. It summarizes molecular similarity in cross-sectional tissue and does not establish temporal progression, ancestry or clonal evolution.

### Malignant tissue is organized as a core with stromal and immune compartments outside

Whole-section neighbourhood enrichment showed non-random spatial organization (Fig. 4a,b). Sixty-nine of 120 cell-type pairs passed correction. CAF-malignant (z=-18.4), malignant- myeloid (z=-10.4) and malignant-stellate (z=-8.7) associations were strongly negative, whereas B-T-cell and endothelial-stellate relationships were positive. Section-resolved summaries showed that these patterns were not carried by one specimen (Fig. 4c). Recurrent cellular neighbourhoods independently summarized normal-exocrine, malignant, stromal and immune local compositions (Fig. 4d,e): the normal-exocrine CN0 fell from 69.3% of Normal spots to 0.75% of PDAC spots, whereas PDAC-associated CN7 and CN9 rose to 29.56% and 17.96%.

**Fig. 4.**
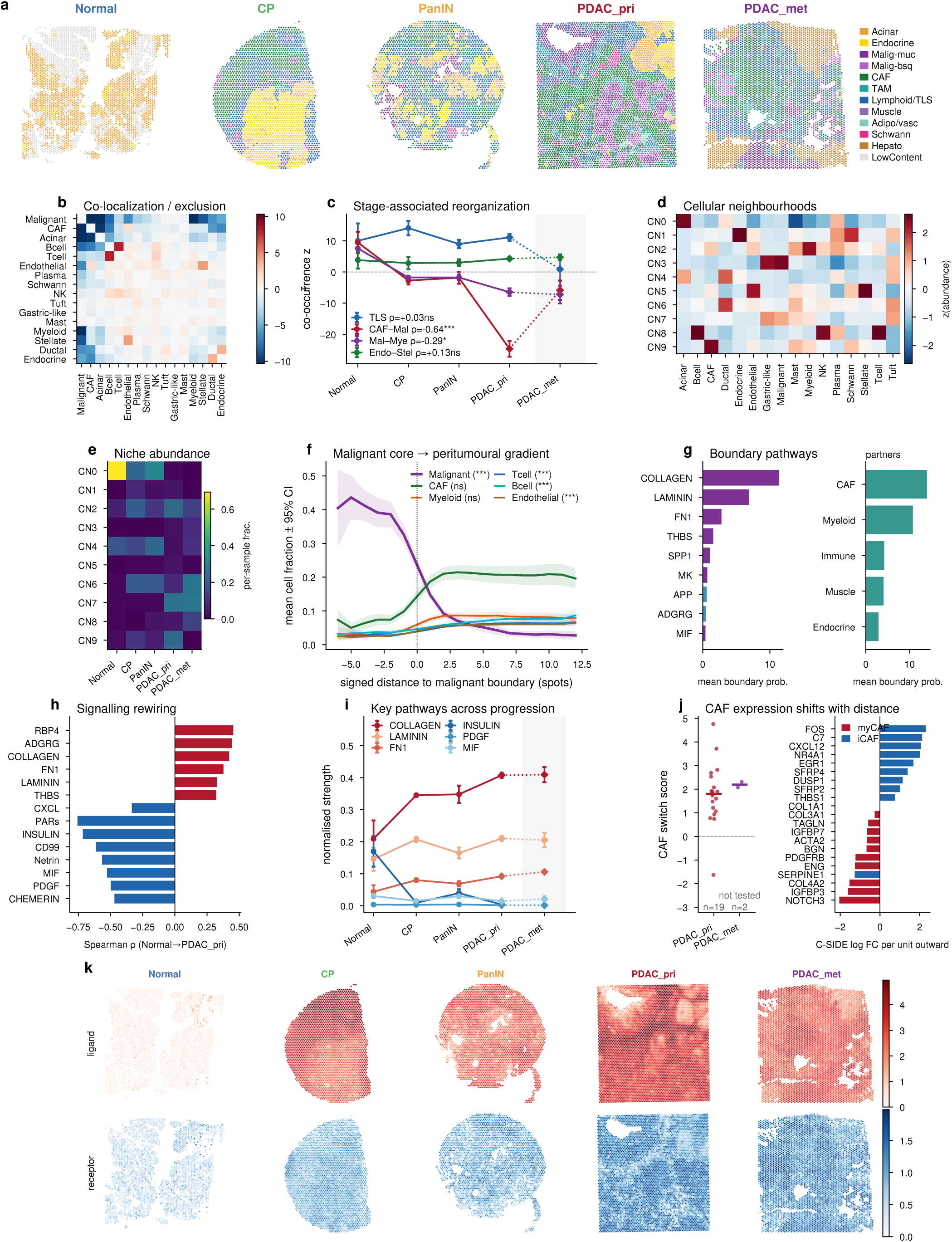
Recurrent spatial compartments and candidate communication at the malignant boundary. **a**, Representative BANKSY maps. **b**, Cohort-wide Squidpy neighbourhood-enrichment z scores; 69 of 120 pairs pass correction. **c**, Priority pairs summarized per section and group. **d**, Recurrent cellular-neighbourhood composition. **e**, Neighbourhood abundance by group. **f**, Mean cell2location fraction versus signed distance in 43 PDAC sections. CAF and myeloid boundary-versus-exterior adjusted P=1.0; T and B cells continue to rise outside. **g**, Candidate spatial CellChat pathways and partner domains; COLLAGEN, LAMININ, FN1 and THBS lead. **h**, Sample-level pathway trends. **i**, Per-sample pathway strengths. **j**, C-SIDE CAF expression versus signed distance in 19 primary-PDAC patients from five studies (one-sided Wilcoxon P=4.8x10^-5); two metastatic patients are displayed but not tested. **k**, Representative COLLAGEN ligand and receptor RNA fields. Communication panels report candidate associations, not direct signalling.

Global segregation can coexist with concentrated interaction at a narrow interface. We therefore measured signed distance from each spot to the malignant-domain boundary in 43 PDAC sections (Fig. 4f). Malignant fraction declined from 0.363 in the interior to 0.040 outside. CAF fraction increased from 0.086 to 0.196 at the boundary and 0.198 outside; myeloid fraction rose from 0.037 to 0.079 and 0.083. For CAFs and myeloid cells, boundary and exterior abundance were indistinguishable after correction (adjusted P=1.0). T- and B-cell fractions continued to rise beyond the boundary. The data therefore support a malignant core, a CAF/myeloid surround beginning at the edge, and a more distal lymphoid compartment. They do not support a universal narrow stromal rim or physical barrier.

Distance-constrained CellChat analysis identified candidate extracellular-matrix relationships at this interface (Fig. 4g). COLLAGEN, LAMININ, FN1 and THBS were the leading pathways, with the strongest partner-domain scores in CAF and myeloid tissue. Along the FFPE disease axis, COLLAGEN, FN1, LAMININ, THBS and ADGRG increased, whereas MIF declined and SPP1 did not pass correction (Fig. 4h,i). These probabilities combine RNA abundance, prior ligand-receptor knowledge and distance; they are candidate associations, not measurements of protein binding or causality.

C-SIDE then separated fibroblast state from fibroblast abundance. Across 19 primary-PDAC patients from five studies, boundary-proximal myofibroblastic genes gave way to inflammatory fibroblast genes farther outside (one-sided Wilcoxon P=4.8x10^-5; Fig. 4j). Two metastatic patients were displayed but not tested. RNA-weighted COLLAGEN ligand and receptor fields localized around representative malignant-stromal interfaces (Fig. 4k). Together, the data connect a graded fibroblast-state transition with an ECM-rich candidate interaction field.

### Six patient-matched pairs reproduce the principal tumour-associated changes

We retained six matched Normal-tumour pairs from two studies, allowing each patient to serve as their own control. Endogenous *S100P*, deconvolved malignant abundance, malignant-domain coverage and EMT changed in the same direction within the pairs (Fig. 5a). *S100P* was 59-fold higher in tumour, and malignant domains occupied a median of 0% of Normal tissue and 32.7% of PDAC tissue. These are correlated views of one tissue change, not four independent validations.

**Fig. 5.**
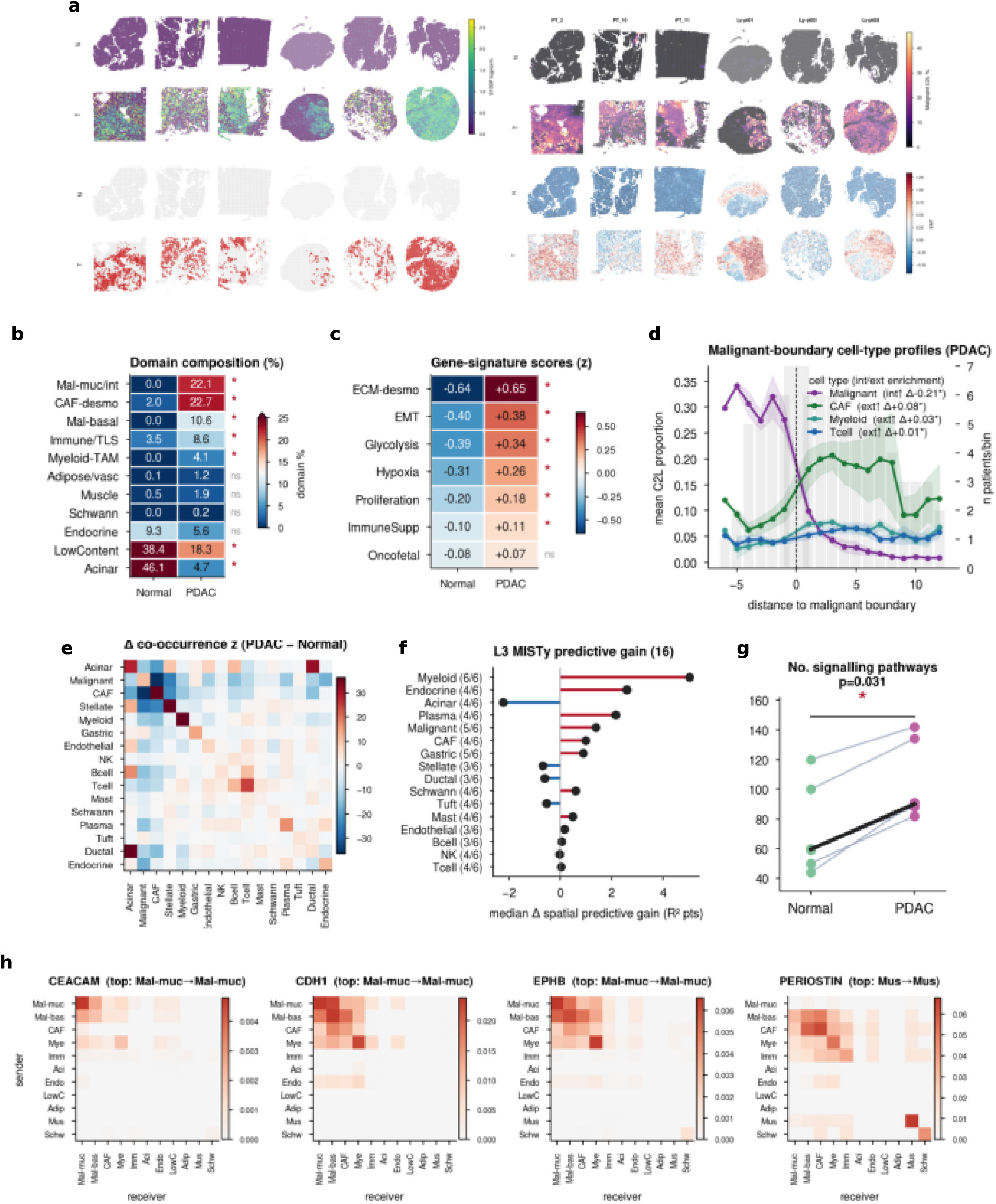
Within-patient confirmation of tumour-associated spatial remodeling. Six matched Normal-tumour pairs were retained across two studies. **a**, Endogenous *S100P*, malignant abundance, malignant-domain membership and EMT. **b**, Paired BANKSY-domain composition with exact patient-level tests. **c**, Paired molecular programs; oncofetal q=0.0625 is not significant. **d**, PDAC signed-distance profiles; asterisks denote 6/6 concordance and exact P=0.03125. **e**, PDAC-minus-Normal pooled neighbourhood-enrichment matrix, shown descriptively. **f**, Paired MISTy gain; myeloid gain rises in 6/6 but has q=0.50. **g**, Retained CellChat pathways per section; median paired increase 30.5, exact P=0.03125. **h**, CEACAM-, CDH1-, EPHB- and PERIOSTIN-associated relationships retained in pooled PDAC. Retained does not mean biologically absent from Normal.

Malignant mucinous/intestinal, basal/squamous, CAF-desmoplastic, immune-lymphoid and myeloid domains increased, whereas acinar and low-content domains declined in all six patients (Fig. 5b). ECM remodeling, EMT, glycolysis, hypoxia, proliferation and immune-suppression scores also rose in all six; oncofetal reactivation did not pass correction (q=0.0625; Fig. 5c). With six pairs, the minimum two-sided exact sign-flip P value is 0.03125, so effect size and directional concordance were emphasized.

The boundary geometry was likewise concordant (Fig. 5d). Exterior-minus-interior differences were -0.214 for malignant cells, +0.080 for CAFs, +0.033 for myeloid cells and +0.015 for T cells, with all six patients sharing each direction. A pooled arm-level co-occurrence difference matrix was descriptive (Fig. 5e). In the paired MISTy analysis, myeloid multiview gain increased in all six by a median 5.03 R2 percentage points, but did not survive correction (q=0.50; Fig. 5f).

The number of CellChat pathways retained after common filters increased in every tumour section by a median of 30.5 (exact P=0.03125; Fig. 5g). CEACAM-, CDH1-, EPHB- and PERIOSTIN-associated relationships were retained in the pooled PDAC arm (Fig. 5h). Retention by the pipeline does not prove biological absence from Normal tissue. The paired analysis therefore supports the main composition, program and boundary changes while leaving communication as a candidate mechanism.

### Visium HD exposes the limits of contact inference from spatial bins

Two public Visium HD sections were segmented into 176,674 Normal and 142,540 PDAC cells. Because the Normal section was FFPE and probe based and the PDAC section was fresh frozen with 3-prime whole-transcriptome chemistry, they were analyzed independently. BANKSY recovered coherent tissue compartments (Fig. 6a,b), and multinomial-logistic transfer from the 17-type reference delineated malignant nests and their boundary in PDAC (Fig. 6c).

**Fig. 6.**
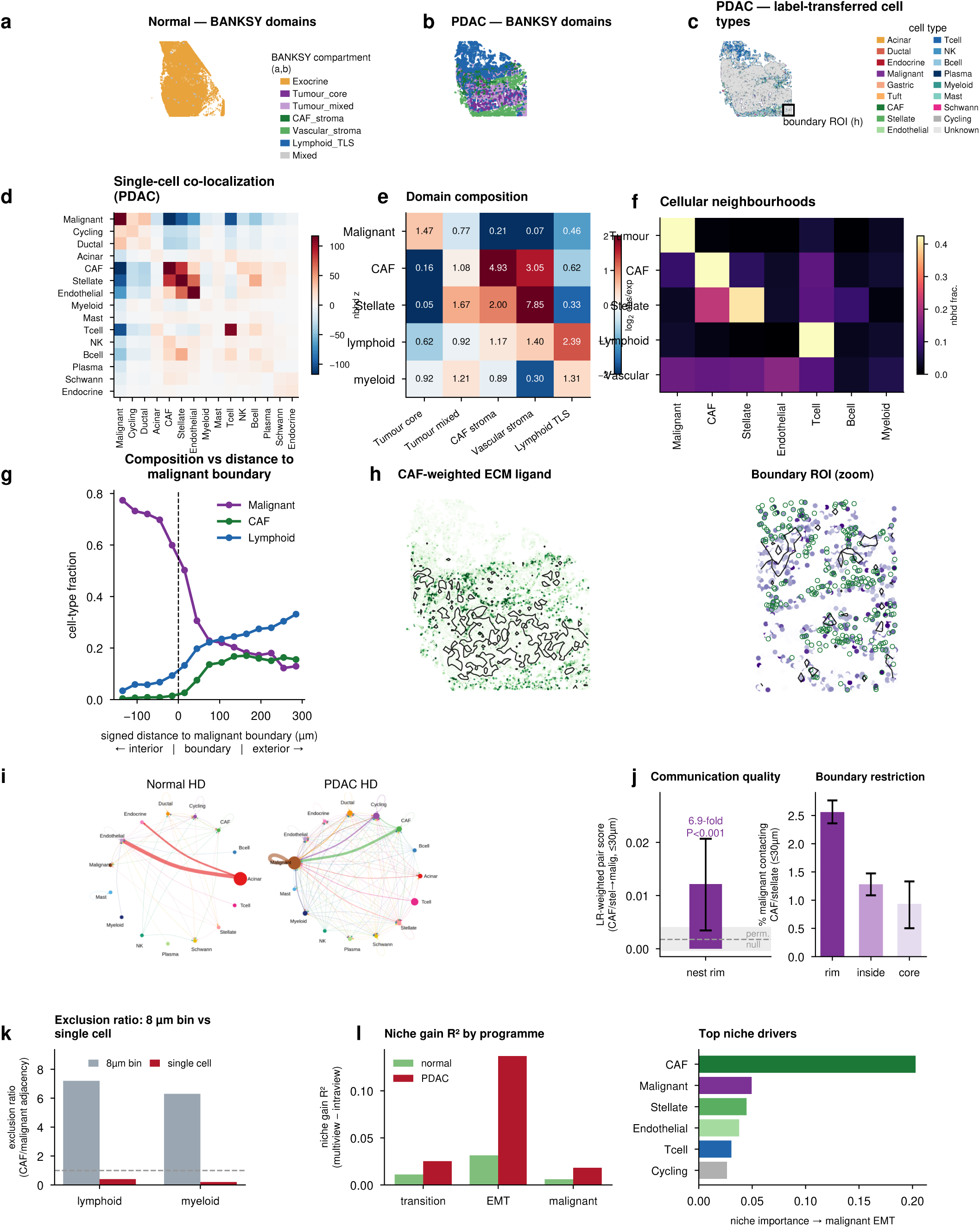
Visium HD tests the spatial architecture at single-cell resolution. The Normal and PDAC sections contain 176,674 and 142,540 segmented cells and differ in preservation and chemistry. **a,b**, BANKSY domains in Normal and PDAC. **c**, Reference-transferred PDAC cell types and boundary region. **d**, Single-cell neighbourhood enrichment. **e**, Observed-to-expected lineage abundance across compartments. **f**, Recurrent cell-scale neighbourhoods. **g**, Cell fractions versus signed distance. **h**, CAF-weighted ECM RNA field. **i**, Aggregate CellChat networks; the cross-chemistry comparison is directional. **j**, CAF/stellate-to-malignant candidate score versus label permutation (6.89-fold, P=4.998x10^-4). **k**, Exterior-to-interior immune ratios from 8-um bins and segmented cells in the same section. **l**, MISTy gain and ranked niche contributors. Effects are within representative sections, not patient-level prevalence estimates.

Single-cell neighbourhood enrichment again showed global malignant-CAF, malignant-T-cell and malignant-stellate segregation (Fig. 6d). Compartment enrichment and recurrent neighbourhoods resolved malignant-core, CAF-stromal, vascular and lymphoid/TLS local states (Fig. 6e,f). Signed-distance profiles placed malignant cells inside and CAF and lymphoid cells outside (Fig. 6g), while a CAF-weighted ECM RNA field localized to the same interface (Fig. 6h).

Aggregate CellChat networks differed directionally between the cross-chemistry sections (Fig. 6i). Within the PDAC section, however, the CAF/stellate-to-malignant candidate score was 6.89- fold above the label-permutation mean (P=4.998x10^-4), and candidate contacts were most frequent in the boundary band (Fig. 6j). These remain RNA- and prior-based associations.

Aggregating the same section into 8-um bins reversed the estimated immune gradient: exterior-to-interior ratios for lymphoid and myeloid compartments were 7.2 and 6.3 in bins but 0.4 and 0.2 after cell segmentation (Fig. 6k). MISTy identified EMT as the program with the largest neighbourhood-associated gain (R2 gain=0.137), with CAFs the top-ranked niche contributor (Fig. 6l). This single section supports cell segmentation for contact-scale questions and shows that small bins should not automatically be treated as cells.

### Xenium recovers recurrent neighbourhoods but sample-specific boundaries

We analyzed one Normal and three PDAC Xenium sections containing 103,859, 122,445, 190,455 and 220,837 cells after quality control. Panel-restricted CellTypist models were trained from the integrated reference. The three PDAC v2 annotations passed prespecified lineage gates, but the Normal classifier recovered only 0.03% T cells despite an independent marker audit indicating approximately 1.4%. Normal lymphoid abundance is therefore displayed as below panel detection and is not used for a Normal-PDAC comparison (Fig. 7a-c).

**Fig. 7.**
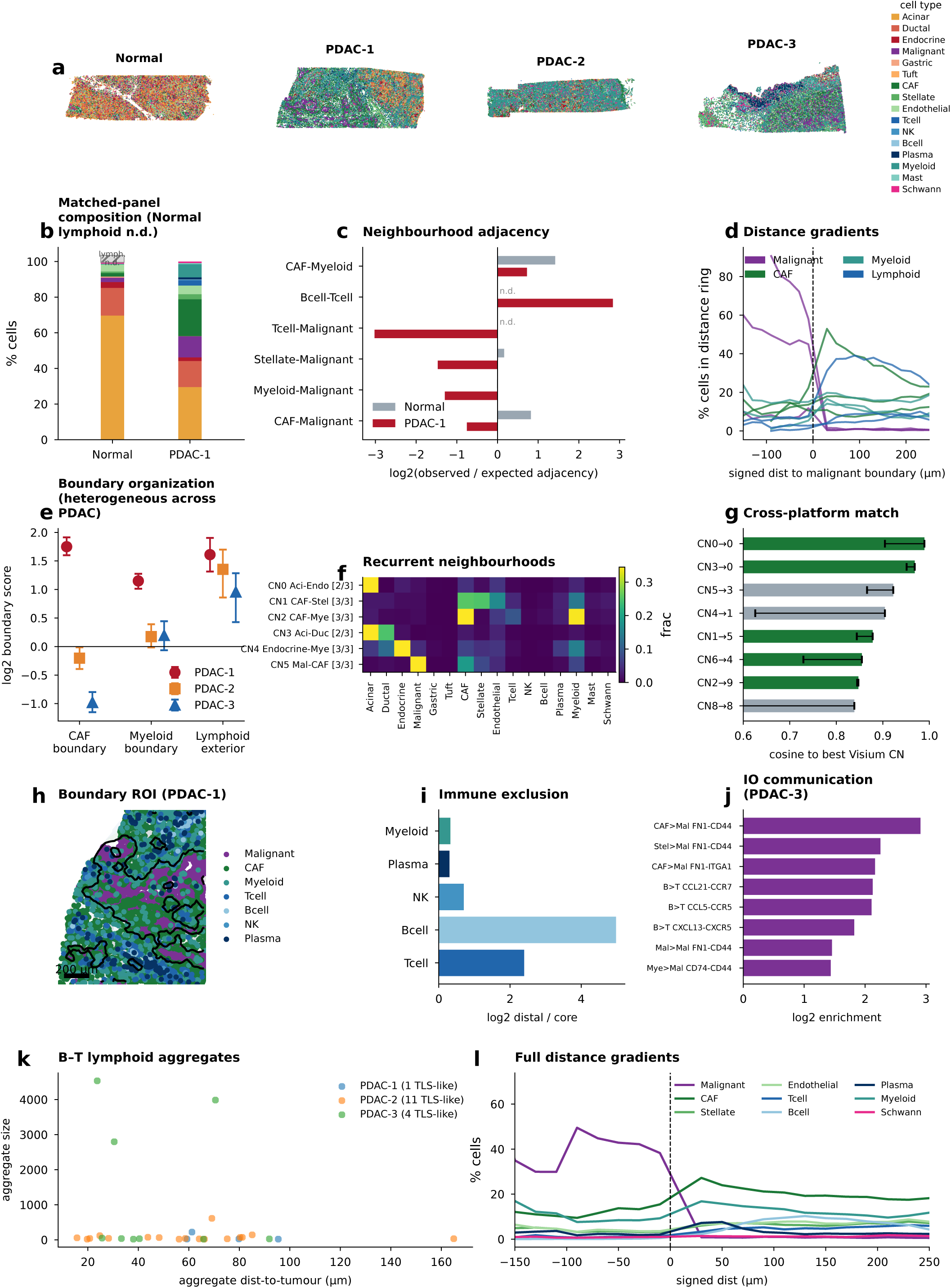
Xenium resolves recurrent and sample-specific cell-scale spatial geometry. One Normal and three PDAC sections contain 103,859, 122,445, 190,455 and 220,837 cells after quality control. **a**, v2 cell-type maps. **b**, Shared-panel Normal and PDAC-1 composition; the hatched Normal segment is lymphoid below panel detection. **c**, Matched-panel observed-to- expected adjacency; Normal lymphoid pairs are not determined. **d**, Signed-distance profiles in three PDAC sections. **e**, CAF boundary, myeloid boundary and lymphoid exterior scores with spatial block-bootstrap intervals. **f**, Recurrent joint cellular neighbourhoods. **g**, Best Xenium-to- Visium composition correspondence among entries with nominal label-permutation P<0.1; green denotes nominal P<0.05 and the left endpoint marks the second-best match. **h**, PDAC-1 boundary region. **i**, Within-PDAC immune distal-to-core ratios. **j**, Significant panel-aware candidates in PDAC-3; complete COLLAGEN, SPP1 and MIF axes are absent, and conventional CellChat retained none. **k**, Operational B-T aggregates. **l**, Full distance gradients. Xenium does not support a Normal-PDAC immune comparison or patient-level prevalence inference.

Within PDAC, signed-distance profiles placed malignant cells inside and lymphoid cells outside, but stromal boundary behavior varied (Fig. 7d,e). The CAF boundary score was 1.75 in PDAC-1, -0.20 in PDAC-2 and -0.98 in PDAC-3. Myeloid boundary enrichment was clear only in PDAC- 1. In contrast, lymphoid-exterior scores were positive in all three sections (1.61, 1.35 and 0.96). Xenium thus supports an outward lymphoid tendency but not a universal CAF or myeloid boundary peak.

Eight of nine joint cellular-neighbourhood classes exceeded 1% abundance in at least two PDAC sections, including CAF-stellate, CAF-myeloid and malignant-CAF classes in all three (Fig. 7f). Five classes matched Visium neighbourhood compositions more strongly than nominal label- permutation controls, although shared reference labels make this correspondence rather than independent validation (Fig. 7g). A PDAC-1 region and immune distal-to-core ratios show the cell-scale geometry directly (Fig. 7h,i).

Targeted-panel coverage imposed a stricter limit on communication. Complete COLLAGEN, SPP1 and MIF axes were absent from every panel, and conventional CellChat retained no significant interaction. A separate panel-aware distance-permutation analysis detected assay- observable candidates in PDAC-3, including 7.51-fold CAF-to-malignant FN1-CD44 enrichment (FDR=0.0216; Fig. 7j). These data cannot validate the full cohort-wide ECM result.

The spatial rule identified 1, 11 and 4 B-T aggregates meeting the operational TLS-like criterion in PDAC-1, PDAC-2 and PDAC-3 (Fig. 7k). Their size and distance varied widely, and targeted RNA panels cannot establish full TLS histology. Full cell-type profiles further showed that the broad core-to-exterior organization was assembled differently in each section (Fig. 7l). Xenium therefore functions as a selective in-situ stress test: recurrent neighbourhoods and lymphoid- exterior geometry remained visible, whereas stromal boundaries were sample specific and key ligand-receptor axes were incompletely observable.

## Discussion

This study connects a large human pancreas single-cell reference to spatial measurements spanning conventional Visium, matched patients, Visium HD and Xenium. Two findings organize the analysis. First, epithelial regions can be placed along a reproducible cross-sectional coordinate extending from acinar-rich to malignant-rich tissue. Second, PDAC commonly contains a malignant core with CAF and myeloid compartments rising from its edge and lymphoid compartments displaced farther outside. The matched analysis shows that these changes persist when each patient serves as their own control, while the higher-resolution assays define where measurement scale and targeted panels alter the result.

The epithelial axis is deliberately not called a lineage trajectory. Five algorithms agreed because the data contain a strong continuous difference between differentiated exocrine and malignant tissue, but they shared the same embedding, root and upstream epithelial labels. Agreement therefore excludes dependence on one trajectory implementation; it does not provide five independent histories of tumour evolution. Cross-sectional sampling also cannot establish whether high-score malignant tissue passed through the intermediate states represented elsewhere. Longitudinal tissue, lineage tracing or matched phylogenetic information would be required for that inference.

The spatial result resolves an apparent contradiction. At whole-section scale, malignant tissue was negatively associated with CAF and myeloid compartments. At the interface, those compartments increased sharply from the malignant interior and remained abundant beyond the boundary. Whole-section neighbourhood enrichment asks whether labels broadly occupy the same tissue, whereas signed distance asks how composition changes along their interface. Both observations can be true. Importantly, neither the 43-section Visium profile nor the three Xenium sections support a universal narrow fibroblast or myeloid rim. The more defensible model is a malignant core with a stromal and myeloid surround beginning at its edge and a lymphoid compartment farther outside.

Extracellular-matrix pathways dominated candidate communication at this interface. COLLAGEN, LAMININ, FN1 and THBS increased along the FFPE disease axis, and C-SIDE identified a shift from boundary-proximal myofibroblastic genes to more external inflammatory fibroblast genes. This links spatial position to fibroblast state rather than number alone. It remains an observational inference. CellChat probabilities, RNA-weighted fields and MISTy predictive gains do not measure protein binding or causal regulation. The Xenium result reinforces this boundary: an FN1-CD44 association was visible where the IO panel allowed it, but complete COLLAGEN, SPP1 and MIF axes could not be tested.

The six matched pairs are the most direct control for public-cohort confounding, although they came from two studies and are not a prospective validation cohort. Their complete directional concordance argues against patient composition and study source fully explaining the main domain, program and boundary changes. The small n constrains attainable significance and motivated reporting effect sizes, concordance and exact tests. The post-hoc exclusion of one Normal section for purity remains a limitation and is documented in the source ledger.

Visium HD and Xenium answer narrower questions. The HD section showed that 8-um aggregation could reverse immune-gradient estimates relative to segmented cells, cautioning against treating small bins as direct contact measurements. Xenium showed recurrent neighbourhood composition but heterogeneous stromal boundary scores. Its Normal classifier failure and non-identical panels prevent a Normal-PDAC immune-abundance comparison. Neither platform supplies patient-level prevalence estimates, and both reuse the integrated reference for biological labels.

Other limitations follow from the public-data design. Metadata and clinical annotations are incomplete, precursor and chronic-pancreatitis groups are smaller than the PDAC group, and whole-section labels can conceal mixed pathology. Marker-defined malignant identity is not genomic proof, but it was preferable to copy-number calls that failed known-diploid controls. The spatial cohort lacks harmonized outcomes, and ligand-receptor analyses remain hypothesis-generating. Several computational layers also share upstream labels; agreement across them is robustness to implementation, not full evidential independence.

The resource is intended to be reusable. Its reference can support deconvolution of new pancreatic spatial datasets, its domain vocabulary can be transferred independently of the epithelial axis, and its sample- and patient-level tables allow each conclusion to be retested as new cohorts arrive. The next decisive step is not simply adding more unmatched sections. Matched longitudinal tissue with histopathology, genomic measurements, treatment information and spatial protein assays will be needed to determine whether this architecture is stable, reversible or predictive of therapeutic response.

## Methods

### Study design and terminology

This secondary analysis used public, de-identified human single-cell, single-nucleus and spatial transcriptomic data. Evidence was organized into a multi-study reference, a standard-Visium discovery cohort, patient-matched Normal-tumour sections and representative Visium HD and Xenium sections. Healthy donor pancreas and tumour-adjacent Normal tissue were retained as distinct source groups. “Control” denotes their combined use only in specified tests. The epithelial axis is transformation associated and cross-sectional; communication denotes a candidate association inferred from RNA, prior ligand-receptor knowledge and geometry.

### Single-cell reference, integration and annotation

Count matrices and author metadata from 19 studies were harmonized to stable cell, study, sample, patient, disease, treatment, assay and library fields. PBMC-derived cells, organoids and treatment-specific observations were excluded from the tissue reference used for the primary PDAC-Control analyses. Newly ingested cells required at least 200 detected genes and mitochondrial fraction no greater than 20%; the curated base atlas retained study-specific quality control. The combined pre-annotation matrix contained 1,187,920 cells and 19,633 genes; removal of doublets, erythroid and excluded cells yielded 1,186,130 cells.

Five thousand highly variable genes were selected from raw counts with the Seurat v3 flavor^35^. Counts were normalized to 10,000 per cell, log1p transformed, scaled with clipping at 10 and reduced to 50 principal components in a Scanpy-compatible workflow^36^. Harmony used sample as the batch variable, a 20-iteration maximum and seed 0 and converged after nine iterations^37^. RAPIDS-singlecell computed a 15-nearest-neighbour graph, Leiden clusterings and UMAP.

scVI was the principal alternative integration^38^. It used the same 5,000 HVGs and raw counts, sample as batch, 30 latent dimensions, two hidden layers, negative-binomial likelihood, a 200- epoch maximum and early stopping. Harmony, scVI and unintegrated PCA were compared on a seeded 100,000-cell subset with scIB metrics. Aggregate scores were 0.6218 for Harmony and 0.6190 for scVI, so Harmony was retained without interpreting the small difference as general superiority.

Broad labels were mapped from the curated atlas, split where mixed clusters required greater resolution and reviewed with canonical markers. Malignant cells required epithelial identity, a tumour-associated marker program and loss of the corresponding normal exocrine program. The final vocabulary contained 17 broad types and 28 states. inferCNV and CopyKAT were run per sample as malignancy audits^39^; their known-diploid failures excluded them from label assignment.

For composition inference, samples had to come from one of five studies with both PDAC and Control arms and lack author-reported sorting or grossly non-representative composition. Cell-type percentages were compared with two-sided Wilcoxon rank-sum tests and Benjamini-Hochberg correction. Expression and signature scores were summarized by sample within cell type. Cells were never treated as independent patient replicates.

### Standard Visium processing, mapping and domains

The standard spatial cohort comprised 176 sections and 458,877 tissue spots from six public sources. Counts, coordinates and sample, study, preservation, treatment and pathological-group metadata were joined by stable identifiers; duplicate representations were removed. Sections remained separate during graph construction. The main Normal-chronic-pancreatitis-PanIN- primary-PDAC trend was restricted to 75 FFPE samples because preservation and disease were partially collinear.

cell2location 0.1.5 mapped 17-type reference signatures to spatial counts and exported posterior fifth-percentile abundance estimates^40^. RCTD/spacexr 2.2.1 provided a second mapping with a 16-type reference in doublet/full mode^41^. Both use the same reference, so concordance was interpreted as cross-algorithm robustness. The production cell2location launch command was not retained; script defaults of 20 expected cells per location, detection alpha 20 and 1,300 epochs are not asserted as recovered production values.

BANKSY used counts normalized to 10,000, log1p, 2,000 sample-aware HVGs and k_geom=6^42^. Lambda was 0.2 for cell typing and 0.8 for tissue domains. The BANKSY matrix was reduced by PCA, integrated with Harmony on sample and clustered at Leiden resolution 0.5. Domain names combined endogenous markers, cell2location and RCTD composition. Low-content or tiny domains remained visible in maps but were excluded from domain-level inference.

Stage-associated expression was aggregated within sample and ranked by Spearman association with the ordered FFPE group. Domain fractions and program scores were likewise tested at sample level with Benjamini-Hochberg correction. Hallmark enrichment used fgsea on the complete ranked list.

### Epithelial axis and spatial boundary analysis

Epithelial spatial units required summed cell2location epithelial weight of at least 0.5. The 83,384 selected spots from 84 FFPE samples were represented by 2,000 HVGs, 50 PCs, sample-level Harmony and a 15-nearest-neighbour graph. Diffusion pseudotime, Slingshot, VIA, Palantir and Monocle3 were rooted in acinar epithelium^43–47^. Oriented ranks were scaled to 0-1 and averaged. Evaluation used pathological-group recovery, cross-tool agreement, Moran’s I, malignant-abundance association, endpoint composition and bootstrap stability. Leave-one-study-out and unintegrated runs were sensitivity analyses. The historical Monocle3 version remains a provenance gap.

Squidpy neighbourhood enrichment was calculated within section with label permutation^48^. For signed-distance analysis, malignant tissue was the union of malignant mucinous/intestinal and basal/squamous BANKSY domains. Sections required at least 30 malignant and 50 non-malignant spots. Distance was normalized by median spot spacing, signed negative inside, and summarized per section in interior, boundary and exterior zones. Mixed-effects models compared zones, with paired Wilcoxon fallback and within-family correction.

Spatial CellChat used CellChatDB.human, population-size weighting, distance use, 250- micrometer interaction range, truncated mean with trim 0.1 and 20 bootstraps^49,50^. Ligand and receptor fields were spatially weighted RNA scores. C-SIDE estimated CAF-specific expression against signed distance while conditioning on other cell-type proportions, and patients were the inferential units. MISTy quantified incremental spatial predictive information^51^. None of these statistics was interpreted as direct signalling or causal regulation.

### Patient-matched analysis

Six matched pairs were retained: Khaliq PT_2, PT_10 and PT_11, and Lyman pt01, pt02 and pt03. Lyman_pt05 was excluded after a post-hoc Normal-purity check. Paired effects used exact sign-flip or signed-rank tests at patient level and Benjamini-Hochberg correction within stated families. Pooled co-occurrence matrices were descriptive. CellChat pathway counts and MISTy gains were calculated per section before pairing.

### Visium HD

Two public 10x Genomics datasets were analyzed: an FFPE probe-based non-diseased pancreas and a fresh-frozen 3-prime whole-transcriptome PDAC, both distributed as Space Ranger 4.0.1 outputs. Their chemistry and preservation differ, so sections were analyzed independently.

StarDist 0.9.2 and bin2cell 0.3.4 assigned counts to segmented cells^52^. BANKSY defined compartments. Cell types were transferred by multinomial logistic regression from up to 8,000 reference cells per type; reference and query were normalized to 10,000 and log transformed, and predictions below confidence 0.5 were Unknown.

Physical-coordinate neighbours defined HD adjacency. The malignant mask and signed-distance bins were derived from transferred malignant cells. The same PDAC section was aggregated into 8-um bins for the resolution comparison. Boundary candidate scores were compared with label permutations and spatial block resampling.

### Xenium v2 analysis

Four public Xenium sections were used^53^: one Normal and one stage III PDAC on a 377-gene panel, one grade I-II PDAC on the 377-gene panel plus 97 add-on genes, and one stage IIB grade 3 PDAC on a 380-gene immuno-oncology panel. Xenium Onboard Analysis versions were 1.5.0, 2.0.0, 1.6.0 and 2.0.0. Cells with fewer than 10 transcripts or five genes and negative-control probes were removed.

Panel-specific CellTypist models were trained from the integrated reference after restriction to each panel, normalization to 10,000 and log transformation^54^. Full multinomial fitting was used without majority-vote smoothing. Predictions retained confidence and marker-audit fields. The three PDAC v2 annotations passed prespecified T-cell-recall and rare-type gates; Normal failed lymphoid recall and was reported as below panel detection.

Adjacency used six physical-coordinate neighbours. Malignant masks were rasterized on a 20- micrometer grid, closed for two iterations and filtered to remove islands smaller than five cells. Signed distance was summarized with prespecified rings and 1,000 resamples of 200-micrometer spatial blocks. Joint cellular neighbourhoods used 15-neighbour composition, centered-log-ratio transformation and k-means clustering into nine classes. Cross-platform correspondence compared cosine similarity of neighbourhood composition with a label-permutation null.

Conventional CellChat and a separate panel-aware distance-permutation statistic were kept distinct; the latter used 200 permutations and FDR correction. B-T aggregates were operational spatial objects and were termed TLS-like only under the stated rule.

### Statistics, software and reproducibility

Tests were two sided unless a directional hypothesis was specified before testing. Benjamini-Hochberg correction was applied within each stated family. Effect sizes, sample counts and direction accompany P values. Cells and spots were nested observations, not biological replicates.

Recorded versions included Scanpy 1.11.5, AnnData 0.11.4, harmonypy 2.0.0, RAPIDS-singlecell 0.15.2, cell2location 0.1.5, RCTD/spacexr 2.2.1, BANKSY 1.2.0, Squidpy 1.8.1, CellChat 2.2.0.9001, mistyR 1.99.12, omicverse 2.2.3, bin2cell 0.3.4, StarDist 0.9.2 and Seurat 4.4.0 for v4 delivery objects. scvi-tools 1.4.2 was used for the scVI benchmark. Per-panel sources and statistical units are listed in panel_source_data_map_v3.csv. Remaining provenance gaps are the production cell2location launch parameters, the historical standard-Visium Space Ranger version and the Monocle3 environment/version.

## Data and code availability

All primary datasets are public. Study-level accessions for the 19 single-cell/single-nucleus studies and the standard-Visium cohort are provided in Supplementary Table 1. The harmonized ledger records study, accession, assay, preservation, disease group, treatment and patient identifiers where supplied by the source publication.

The two Visium HD and four Xenium datasets are public 10x Genomics demonstration datasets released under CC BY 4.0. They can be located in the 10x Genomics dataset portal using the identifiers visium-hd-cytassist-gene-expression-libraries-human-pancreas-4, visium-hd-three-prime- human-pancreatic-cancer-fresh-frozen, human-pancreas-preview-data-xenium-human-multi-tissue-and-cancer-panel-1-standard, ffpe-human-pancreas-with-xenium-multimodal-cell-segmentation-1-standard, pancreatic-cancer-with-xenium-human-multi-tissue-and-cancer-panel-1-standard and ffpe-human-ductal-adenocarcinoma-data-with-human-immuno-oncology-profiling-panel-1-standard.

Analysis code, processed metadata, panel-level source tables and environment records are organized in the project release package. A public repository URL or archival DOI has not yet been assigned; the materials are available from the corresponding author upon reasonable request during review and will be deposited in a versioned public repository before journal publication. Raw source matrices remain in their original repositories and are not redistributed where source terms prohibit it. Xenium v2 synchronization and key-file hashes are recorded in pdac_xenium/results/V2_PROVENANCE.md.

## Author contributions

K.L.: Conceptualization, Data curation, Formal analysis, Investigation, Methodology, Software, Validation, Visualization, Writing - original draft, Writing - review and editing. H.Z. and R.S.E.: Data curation, Formal analysis, Investigation, Methodology, Software, Validation, Visualization. Y.S., L.W., A.P.W., T.I.V., C.F., J.P.A. and A.R.H.: Discussion and interpretation of results, Writing - review and editing. W.R.C.: Conceptualization, Funding acquisition, Project administration, Supervision, Writing - review and editing.

## Ethics statement

This study reanalyzed public, de-identified datasets and collected no new human tissue. Ethical approvals and consent procedures are described in the original studies.

